# Flexible parental care in a songbird correlates with sex-specific responses to seasonal phenology, mating opportunity and reproductive success

**DOI:** 10.1101/2025.11.08.687196

**Authors:** Jia Zheng, Wenni Jiang, Hui Wang, De Chen, Zhengwang Zhang, Jan Komdeur

**Affiliations:** Ministry of Education Key Laboratory for Biodiversity Sciences and Ecological Engineering, College of Life Sciences, Beijing Normal University, Beijing 100875, China; Behavioral and Physiological Ecology, Groningen Institute for Evolutionary Life Sciences, University of Groningen, 9712 CP Groningen, The Netherlands; Institute of Ecology and Evolution, University of Bern, Bern, Switzerland

**Keywords:** parental care variations, socio-ecological conditions, mating opportunity, season length, reproductive success, penduline tit

## Abstract

1. Species with diverse parental care patterns often exhibit flexibility in response to environmental changes. Such changes can influence parental care decisions by altering the trade-offs between current and future reproduction. While previous studies have revealed correlations between specific social and ecological factors and parental care patterns, the joint effects of multiple factors could be rather complex and may result in different outcomes within and across populations, and between sexes.
2. In this study, we revealed parental care systems and their seasonal variations across three geographic populations in a polygamous species, the Chinese penduline tits (*Remiz consobrinus*), by monitoring breeding events over years. To investigate how local conditions (including social and ecological conditions) correlate with sex-specific parental care patterns, we compared breeding phenology and local conditions (including season length, mating opportunity and reproductive success) across the populations.
3. Striking population differences in parental care strategies are found in this species: biparental care predominated in one population, whereas female-only care was more common in the other two. Our analyses reveal that, despite a large difference in season length, the time of egg-laying peaked similarly across the three populations. Female penduline tits determined the breeding phenology, which varies male mating opportunities dramatically over the season. For all populations, male Chinese penduline tits were more likely to desert the clutch when male mating opportunities were high. In contrast, females consistently provided care, regardless of variation in female mating opportunities.
4. We also found that fitness costs on the current brood resulting from offspring desertion differed across populations. This desertion cost on brood fitness matters more than achieving a high reproductive reward in driving the emergence of uniparental care, contrary to the theory that uniparental care only occurs when single parents can efficiently raise nestlings alone.
5. Our study demonstrates that similar parental care patterns across populations could result from different underlying factor interactions and highlights the consistent sex-specific responses to local conditions in parental care.

## Introduction

Animals often face reproductive challenges due to environmental changes, such as shifts in phenology caused by global warming or increased predation risks following species invasion (Ryding et al., 2021; Batabyal et al., 2022). To maximize reproductive fitness during environmental changes, individuals need to flexibly adjust their phenotypic traits and reproductive strategies (Miranda, 2017; Ben Cohen & Dor, 2018). Among these strategies, parental care is one of the most plastic behavioural traits (Remeš et al., 2015; Furness & Capellini, 2019). In birds, offspring survival relies heavily on parental provisioning and protection, resulting in biparental care as the main parental strategy (Cockburn, 2006; Remeš et al., 2015). Even so, in a minority of species, uniparental care occurs when one parent deserts its mate and offspring during a specific reproductive stage (often during the nestling phase) to seek additional mating opportunities and thereby potentially increase reproductive output (Roulin, 2002; Griggio & Pilastro, 2007; Ledwoń & Neubauer, 2017; Zheng et al., 2018; Kupán et al., 2021).

This desertion behaviour is conditional on the local environment and may vary considerably both among broods within populations and across breeding seasons (Pilastro et al., 2001; Kupán et al., 2021). Previous studies exploring relationships between environmental conditions and parental care strategies have generally considered single-factor scenarios, comparing care patterns across species/populations experiencing different environmental conditions. For instance, a comparative analysis of six shorebird populations revealed that a skewed adult sex ratio was related to the prevalence of uniparental care, as the rarer sex was more likely to desert the brood and pursue additional breeding to maximize its own fitness (Eberhart-Phillips et al., 2018).

However, the causes of variation in parental care across populations are likely more complex and are attributed to multiple interacting factors and ecological processes. First, different factors, such as skewed adult sex ratio and food scarcity, may exert opposing effects on parental care strategies and jointly influence the distribution of care patterns, potentially leading to misleading conclusions if only a single factor is considered (Balshine-Earn & Earn, 2012; Furness & Capellini, 2019; Vági et al., 2024). Second, recent studies suggest that behavioural responses in parental care can be rather flexible at a temporal scale. For instance, in four boreal owl species (*Glaucidium passerinum, Aegolius funereus, Strix aluco and S. uralensis*) and penduline tits *Remiz spp*., parental care patterns can shift from uniparental to biparental care as the season progresses, driven by the declining fitness returns due to reduced re-mating opportunities or food availability (Pogány et al., 2008; Lehikoinen et al., 2011; Zheng et al., 2021). Similarly, female snowy plovers (*Charadrius nivosus*) adjust their care decisions through desertion by evaluating their partner’s ability to independently complete the remaining nestling-rearing duty (Kupán et al., 2021). Such dynamic environmental conditions within a breeding season may compel individuals to continuously adjust their reproductive decisions in response to concurrent changes in fitness payoffs (Lehikoinen et al., 2011; Béziers & Roulin, 2016). Third, sex-specific roles shaped by selective pressure or physiological constraints in reproduction can intermediate male and female parents respond differently to environmental changes, and lead to variations in parental care system (Donelan & Trussell, 2020; Wang et al., 2023). However, it remains largely unknown how different environmental conditions and their temporal variation shape parental care decisions, and whether these environmental dynamics affect differently on the two sexes in wild populations.

The penduline tit (*Remiz. spp.*) is a small passerine genus renowned for its diverse parental care systems. Among them, the Chinese penduline tits (*R. consobrinus*), distributed across East Asia, exhibit one of the most intricate parental care system: female-only care, male-only care, biparental care and biparental desertion nests all exist within a single population (Ball et al., 2017; Zheng et al., 2021). In addition, Chinese penduline tits exhibit seasonal shifts in parental care, varying from uniparental-female care to biparental care over the course of breeding season within a population (Zhen et al., 2021); the predominant care patterns also observed differed across populations (Zheng, 2022). This complexity is exceedingly rare among passerine birds and presents a great opportunity to investigate how local conditions may shape parental care variations.

In this study, we first revealed the divergent parental care systems in three populations Xianghai (XH), Liaohekou (LHK) and Yellow river delta (YRD) (Figure 1), spanning a geographic range in northeastern China, which is the species’ main breeding area. To explore the potential causes of the variation in parental care, we first compared the breeding phenology and seasonality in parental care patterns across populations. Second, we investigated three key factors that may simultaneously contribute to the variation in the parental care system: (i) mating opportunity, a social condition reflecting the number of available mates, which may temporally vary within populations. Sex-specific offspring desertion can strongly affect mating opportunity as the deserters massively return to the mating pool, thereby affecting future reproductive fitness. (ii) fledging success, a direct measure of current reproductive fitness, reflecting the ability of parent(s) to raise a brood. This success is influenced by a variety of local ecological factors, such as weather conditions, food availability, or predation risks. (iii) season length, the duration of the breeding window available to parents. Its impacts on parental care are complex (Halupka et al., 2021; Zheng et al., 2023) and have received little attention in studies of wild populations (but see Griggio, 2015; Jankowiak & Wysocki, 2016). We predict that brood desertion is more likely to occur in populations with sex-biased mating opportunities and high reproductive success. Season length, which determines the maximal number of breeding attempts, is expected to influence parental care strategy by interacting with the dynamics of mating opportunities. Lastly, previous studies of Chinese penduline tits often observed higher female propensity of care than males (Pogány et al., 2008; Zheng et al., 2021; Zheng, 2022), suggesting that the two sexes may decide on parental care by reacting differently to environmental dynamics.

**Figure 1.**
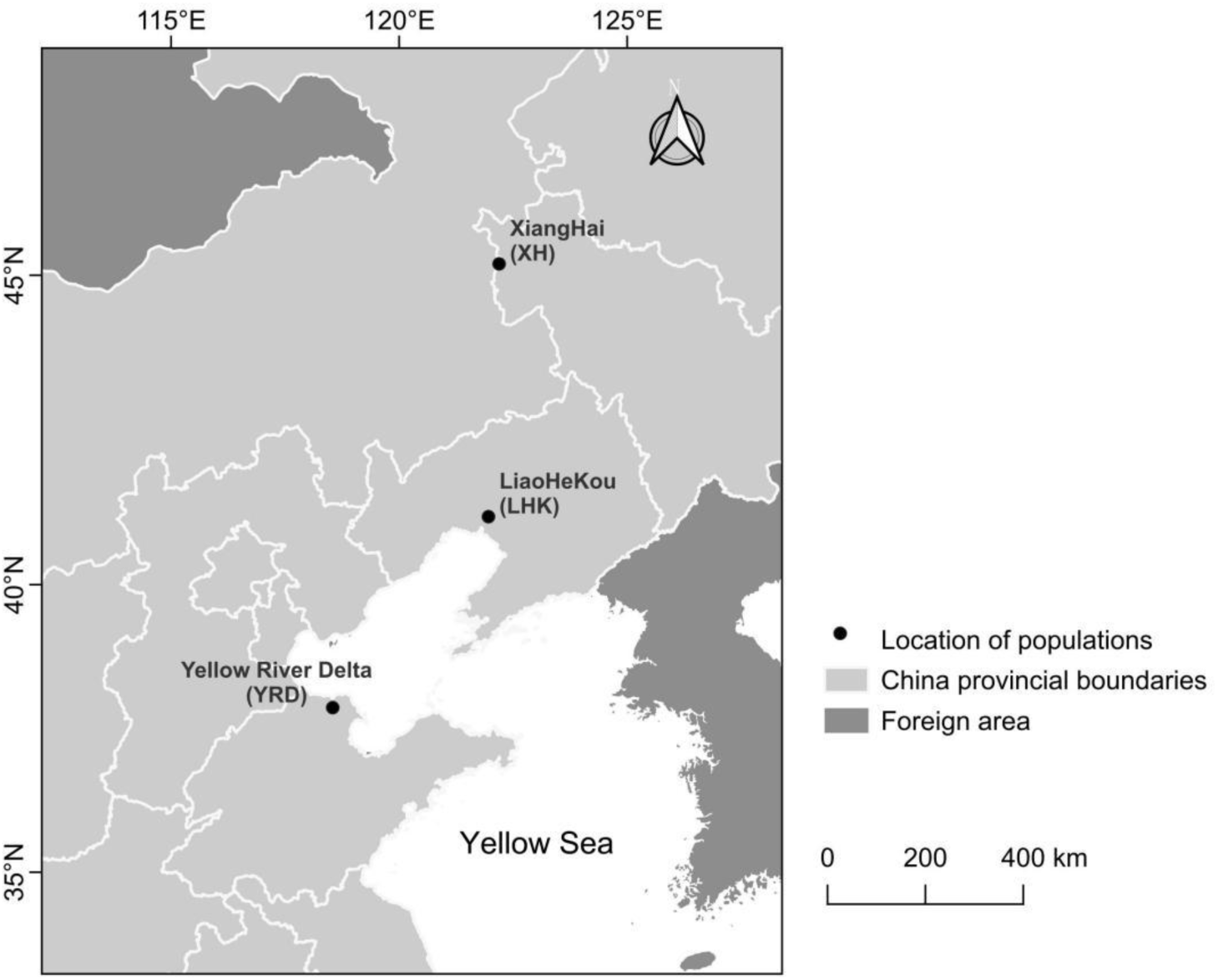
The geographic locations of the three investigated populations of Chinese penduline tits.

## Methods

### Study areas and populations

Chinese penduline tits were studied from 2016 to 2023 in three populations: Liaohekou national nature reserve, Liaoning province, northeast China (LHK population, 40°45’-41°05’N, 121°28’-121°58’ E, 44 km^2^), Xianghai national nature reserve, Jilin province, northeast China (XH population, 44°55’-45°09’N, 122°05’ - 122°31’ E, 15 km^2^) and Yellow river delta national nature reserve, Shandong province, east China (YRD population, 37°45 ’ - 37°52 ’ N, 118°38 ’ – 118°47 ’ E, 17 km^2^, Figure 1). The distance between the LHK and XH population is approximately 530km, and YRD is approximately 700 km south of the LHK population (Figure 1). The migratory Chinese penduline tits arrive in these areas (in March for the YRD population and in late April for the LHK and XH populations) to commence breeding, which lasts until August. Therefore, fieldwork in the LHK population was carried out from May 20th to July 20th in 2016, May 1st to August 1st in 2017 and 2019 (Zheng et al., 2018; Zheng et al., 2021); in the XH population from May 3rd to August 2nd in 2019 and April 25th to August 10th in 2020 (Zheng et al., 2024), and in the YRD population from March 15th to August 20th in 2022 and 2023.

The XH population bred the most northern, that situated in an inland area, surrounded by two endorheic lakes. Meadows cover the Khorchin sandy lands with rolling dunes across the landscape. Trees (large-fruited elm *Ulmus macrocarpa*, Siberian elm, willow *Periploca sepium* and Chinese white poplar *Populus tomentosa*) are scattered across the plain meadows. The LHK breeding site is located in a delta area close to the north end of Bohai Bay. The whole area is covered by natural reed beds, which are traversed by several roads that separate the whole area into nine reed ponds. Trees (Siberian elm *Ulmus pumila*, black locust *Robinia pseudoacacia*, weeping willow *Salix babylonica*, and Chinese white poplar) are intermittently distributed along both sides of the roads and directly next to the water edge of the reed ponds. The YRD breeding site is located at the south end of Bohai Bay. This area comprises a vast expanse of farmland, with several adjacent patches of woods. Water ditches, where reeds grew, traversed between the farmland and woods. Siberian elm, black locust, willow and Chinese white poplar were the main species in the woods, intersected by a few trails.

### Data collection

A few days after arriving at the breeding sites, male Chinese penduline tits initiated nest building at the end of an outside tree branch, meanwhile attracting females with singing and displaying around the nest. A female joins building the nest once it is paired up with a male. Throughout the whole breeding season, we searched the entire study area for new nests on a daily basis by inspecting tree branches or by localizing males through their songs, or tracking their flight routes. When a new nest was found, its nest stage and the tree species used for building were recorded. We divided the nest-building process into six stages (see Figure 1 in Zheng et al., 2018). The time (in days) spent on each stage was recorded by checking the nest every other day. The onset date of nest construction was extrapolated from the nest stage on the day the nest was found. We aimed to discover a nest before the nest stage C (see Zheng et al., 2018, for the process of nest shaping over stages) in order to derive a more precise nest initiation day. Every other day, we made an observation (no longer than 15 min) of each nest that is sufficient to record the parents’ presence, nest-building and mating behaviour and the progress of nest stage using binoculars, staying at 10 m distance from the nest to ensure the breeding adults were not disturbed (Szentirmai et al., 2005; Zheng et al., 2018). A male was defined as paired with a female when it was seen copulating near the nest, and/or staying and weaving the nest together with the female (Szentirmai et al., 2007; Zheng et al., 2018). Before the start of egg laying, males were caught using mist-nets with a dummy male and playback of male songs placed close to the net (shorter than 50 cm), whereas females were trapped in the nest after nestlings hatched with a special tuck net (Zheng et al., 2021). These procedures did not result in nest abandoning. We did not manage to catch all the birds, because we were unable to lure every male in the mist-net and some females could not be caught because the nest was at an inaccessible location (e.g. too high up in the tree). Adults and nestlings were uniquely banded with a numbered metal ring and three coloured rings to distinguish individuals. By doing this, we intended to track every individual’s mating decisions and mating opportunities at the breeding site throughout the entire season (see the section below).

We inspected the high nests (above 3 m) using a 4-m ladder. However, there were some (26%) nests that we could not reach because they were too high (above 5 m) or positioned above the water surface, the parental care pattern could still be scored by observing the parents’ behaviour during the nestling-feeding stage. We calculated the onset of egg-laying based on the number of eggs found in the nests (this species lays one egg every day). Clutch size was determined 10 days after the start of egg-laying (clutch size ranged from 5 to 9 eggs). From the 5^th^ day of egg-laying onwards (with day 1 as laying the first egg), we started to observe the parental care pattern of the nests by daily nest checking, because clutch desertion occurs at the onset of incubation (Zheng et al., 2018). If one sex did not show up during the nest monitoring, a SONY HDR-XR160E video camera was used to film the nest for 2 h continuously. The nest was scored as deserted when one or both parents did not return to the nest during the video recording (Zheng et al., 2021). Since Chinese penduline tits are sexually dimorphic and the sexes can be distinguished by their plumage, we were not only able to monitor the exact date of desertion but also whether the nest was deserted by the male, female, or both sexes.

The hatching date of the first egg was recorded as age 0 and we counted the total number of hatched nestlings 2 days later (all of the nestlings normally hatched within 3 days). Nestlings were ringed with a metal ring on day 5 and individually colour-ringed on day 15 (approximately 6 days before fledging). The number of nestlings on day 15 was considered to be the number of young fledged (Kingma et al., 2008; Zheng et al., 2018). Once a nest failed (i.e., nest vanished, eggs disappeared, nestlings dead or vanished), we recorded the date of nest failure.

### Statistical analyses

Statistical analyses were performed using R version 4.4.3 with R Studio 2024.12.1, and the null hypotheses were rejected at p < 0.05. Mean ± SE was provided in the results. ‘MASS’, ‘dplyr’, ‘car’, ‘lme4’, ‘ggplot2’, and ‘emmeans’ were the main statistical packages used for the analyses. We performed the following analyses across the three populations:

#### Seasonality of parental care

A chi-square test was used to compare the distributions of different parental care patterns between the three populations. A GLM with a logistic distribution was created using the two parental care types (‘uniparental’= 1 and ‘biparental’= 0) as the response variable, nest initiation date, populations, their interactions and year as fixed factors to test the seasonality of the occurrence of different parental care types in the three populations. Year has no effect on either the proportion distributions (*χ_LHK_^2^* = 3.73, *p_LHK_*= 0.71, *χ_XH_^2^*= 2.66, *p_XH_* = 0.44; *χ_LHK_^2^*= 5.36, *p_YRD_* = 0.15) or seasonal variations (*χ^2^*= 0.53, *p* = 0.22) of parental care patterns in the three populations.

#### Season length and breeding timing

To compare breeding phenology of the three populations, we standardized March 10th as the first day for all populations. To test for a difference between the seasonal timing of nest initiations of the three populations, generalized linear models (GLMs) with a Poisson distribution were used. Nest initiation day was the response variable and year and location were the predictors.

#### Breeding timing and parental care

Two GLMs were created to analyze (1) the difference of the first egg-laying day (the response variable) between the three populations (the predictor) with a quasi-Poisson distribution; (2) the difference of clutch size between parental care types among populations with a quasi-Poisson distribution.

Location, care type and their interactions were used as the predictors. Year has no effect on either the first egg-laying day (χ^2^ = 2.55, p = 0.12) or the clutch size (χ^2^ = 0.39, p =0.53), and was therefore excluded from the models.

#### Male mating opportunities and parental care

Mating opportunity for each sex refers to the number of single individuals of the opposite sex that vary over the season. Given the inconspicuous behaviour of female penduline tits, it was impossible to directly estimate male mating opportunity by counting the number of single females in the population. Therefore, we estimated mating opportunities for males by using two measures applied in Zheng et al. (2021): (1) ‘male pairing success’ refers to the possibility that a single male acquires a mate on a season day; (2) ‘male mating time’ refers to the number of days between the day a male initiate the nest and the day it paired up with a female or abandoned the nest due to the failure in mate acquisition. GLMs were used to analyse the seasonal variation of male pairing success over the season with logistic regression, and that of male mating time using Poisson regression. Male pairing success (‘paired’= 1 and ‘unpaired’= 0) and male mating time were the response variables, respectively; nest initiation day, population, their interaction and year were the predictors. As single males always sang and built nests during mate attraction, we estimated the seasonal dynamics of female mating opportunities by summing the number of single males available in the population on every season day.

We then analysed the associations between mating opportunities (for males, using mating time as the response variable; for females, using the number of single males as the response variable) with parental care patterns (i.e., biparental care and uniparental care, as the predictor) by creating GLMs using Poisson distribution. Year has no effect on male pairing success (χ^2^ = 0.08, p = 0.14) and male mating time (χ^2^ = 0.87, p = 0.35) and number of single males (χ^2^ = 0.49, p = 0.22), thereby was not included in the models.

#### Reproductive success and parental care

As measures of reproductive success for each nest with a clutch, hatching success was calculated as the number of hatchlings divided by the clutch size, and fledging success as the number of fledglings divided by the number of hatchlings (Zheng et al., 2018). GLMs with a binomial error were built with (1) the number of hatchlings and unhatched eggs for each nest, and (2) the number of fledglings and the number of dead nestlings of each nest as the response variables. Population ID, parental care strategy and their interaction were taken as the predictors. Year has no effect on hatching success (χ^2^ = 2.70, p = 0.10) and fledging success (χ^2^ = 0.29, p = 0.59), and was therefore excluded from the final models. The differences in hatching success between populations and the differences in fledging success between parental care patterns were analysed using post-hoc tests applied in the two models.

## Results

### Different parental care systems and seasonality across populations

The three populations exhibited distinctive parental care systems (*χ^2^*= 102.23, *df*= 6, *p*< 0.001, Table 1A). Female-only care was predominant in both the LHK and YRD populations, and accounted for more than 70% of the nests. In contrast, biparental care was the dominant pattern (60%) in the XH population (Table 1A), and differed significantly from the other two populations (Table 1B). Notably, biparental desertion was rare in the XH and YRD populations, but more common in LHK (*χ^2^*= 11.06, *df*= 3, *p*= 0.01; Table 1A). Male-only care accounted for small proportions in all three populations. Seasonal trends in parental care varied among populations (*χ^2^*= 17.55, *df*= 1, *p*< 0.001; Figure 2). The LHK population exhibited a clear seasonal shift, transitioning from female-only care to biparental care over the season. Such transition was absent in XH and YRD populations (Figure 2).

**Figure 2.**
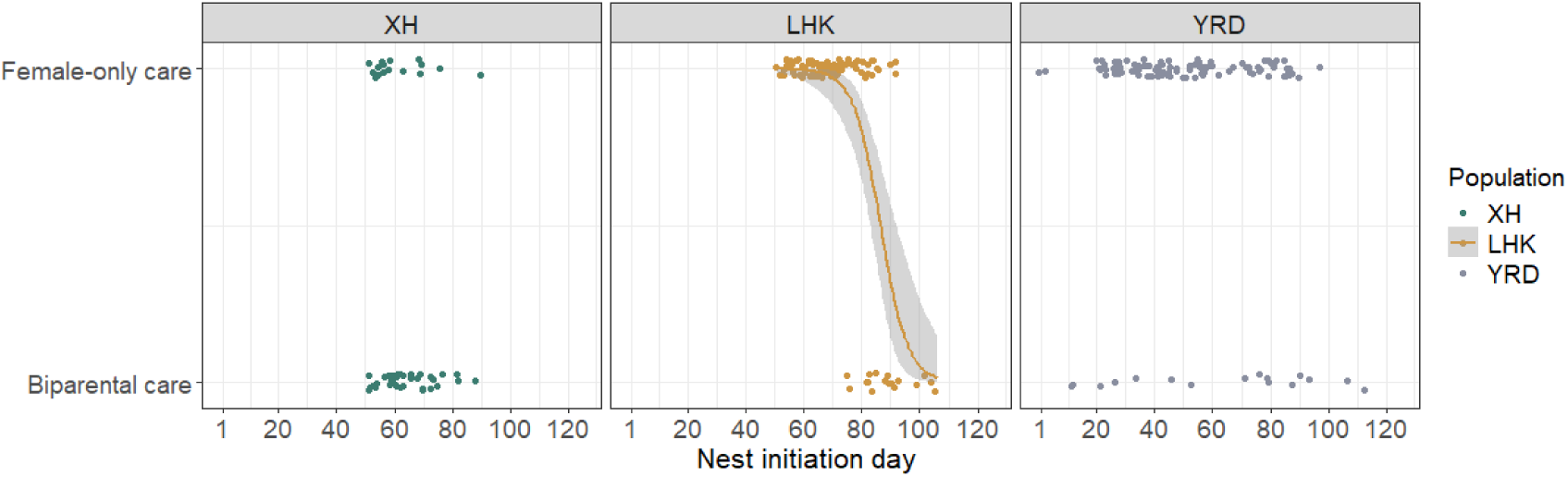
The initiation day of uniparental and biparental care nests over the breeding season in the three populations of Chinese penduline tits. The seasonal trend of parental care variation in LHK population is depicted by logistic regressions, with gray ribbons indicating the 95% confidence level interval. No seasonal trend was found in the XH and YRD populations. *XH: χ^2^*= 0.81, *df*= 1, *p=* 0.38, n = 65 nests; *LHK: χ^2^*= 46.58, *df*= 1, *p<* **0.001,** n= 111 nests; *YRD: χ^2^*= 2.54, *df*= 1, *p=* 0.11, n = 114 nests. X-axis: 1 = March 10th.

**Table 1.** Proportion of parental care patterns in the three populations of Chinese penduline tits. n_LHK_= 154 nests, n_XH_ = 88 nests, n_YRD_ = 128 nests.

| (A) Population | Female-only care | Male-only care | Biparental care | Biparental desertion |
| --- | --- | --- | --- | --- |
| LHK | 71% | 5% | 12% | 12% |
| XH | 31% | 8% | 60% | 1% |
| YRD | 80% | 5% | 13% | 2% |
| (B) | Chi-square ($\chi^2$) | P value | | |
| XH vs LHK | 67.82 | **<0.001** |  |  |
| XH vs YRD | 57.94 | **<0.001** |  |  |
| LHK vs YRD | 11.06 | **0.01** |  |  |

### Difference in nest initiation trends and similar egg-laying peaks across populations

Breeding phenology was similar for each population across years, but varied considerably across the three populations (Table 2, Figure 3). Nest initiation began earliest in the southern YRD population, starting in mid-March, which was about one and a half months earlier than in the other two populations (Table 2). Despite this early start, the last egg was laid around mid-July in all three populations (Table 2), resulting in the YRD population having the longest breeding period, which allowed for two distinct breeding peaks. In contrast, the shorter breeding seasons in XH and LHK populations led to more synchronized nest initiation among males, resulting in higher breeding peaks (Figure 3A). The LHK population exhibited two closely spaced breeding peaks, whereas the XH population had a single but more intensive peak early in the season (Figure 3A).

**Figure 3.**
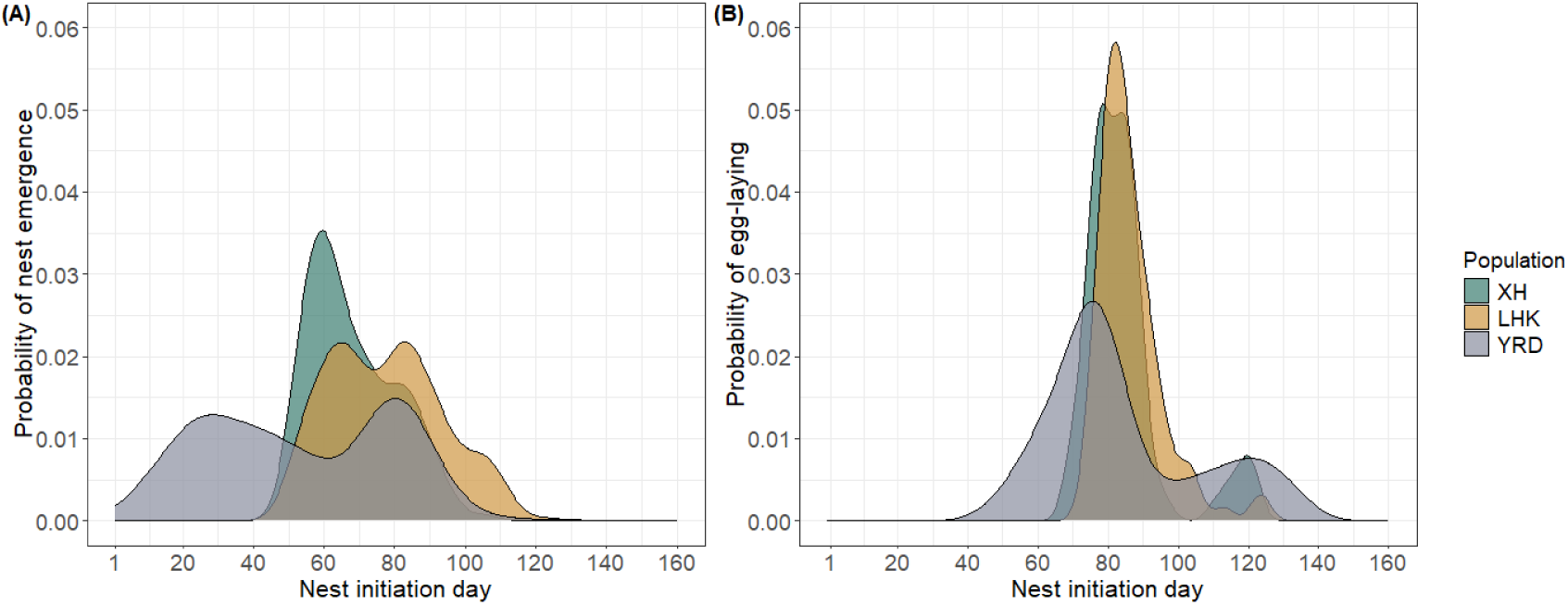
Probability of (A) nest emergence and (B) start of egg-laying day over the breeding season in the three populations of Chinese penduline tits. The total area under a probability density curve is 1, representing the entire probability distribution. The trends of nest emergence were different between the three populations (*χ^2^*= 9.01, *df*= 1, *p*< 0.01, n_XH_ = 244 nests, n_LHK_ = 249 nests, n_YRD_ = 337 nests), The trends of egg-laying were consistent over populations (*χ^2^*= 3.00, *df*= 2, *p*= 0.22, n_XH_ = 107 nests, n_LHK_= 94 nests, n_YRD_ = 143 nests). X-axis: 1 = March 10th.

**Table 2.**
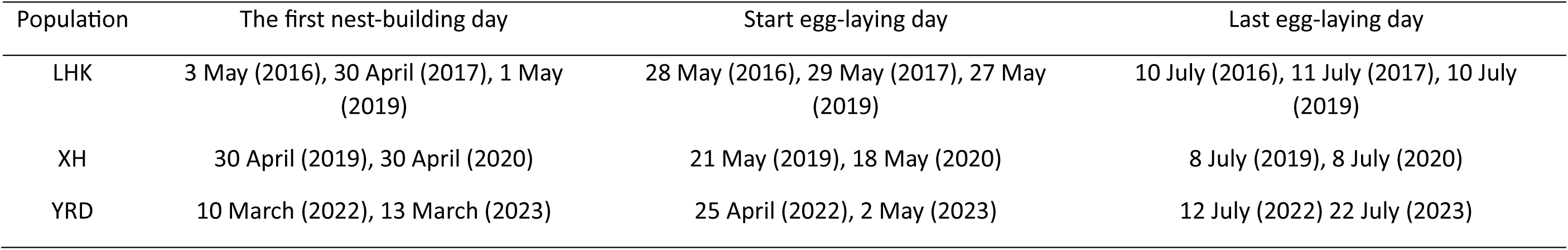
The first day of nest initiation and egg-laying in the three populations of Chinese penduline tits.

In contrast, the timing of egg-laying was similar for each population with a maximum difference of 7 days across years (Table 2). The timing of egg laying also showed less differentiation among populations compared to the first day of nest-building (Table 2, Figure 3b). The first clutch in the YRD population was laid about half a month earlier than in XH and roughly one month earlier than in LHK (Table 2). Despite differences in nest initiation and the timing of first egg laid, the peak of egg-laying was largely synchronized across populations, clustering around days 75-85 (Figure 3b). The extended breeding season in the YRD population showed, in general, a lower peak compared to the other two populations (Figure 3B). Average clutch size differed between the three populations (clutch size mean ± se: LHK--6.80 ± 0.07, n = 94 nests; XH--7.07± 0.08, n= 73 nests; YRD--6.49± 0.06s, n= 143 nests; *F*= 11.62, *df*= 2, *p<* 0.001), but was similar between each of the four care types within each population (*F*= 2.18, *df*= 3, *p*= 0.10).

### Interaction between male mating opportunities and season length and their associations with parental care

Male mating opportunities showed different season trends across the three populations (Figure 4, Table 3). In the LHK and XH populations, male pairing success declined significantly over the season, whereas no such trend was observed in the YRD population (Figure 4A, Table 3A). However, males in the YRD population experienced an ‘efficient pairing window’ around days 55 to 70, during which they achieved high pairing success (24 out of the 27 males paired) and spent shorter mating time (9.2 ± 8.4 days) than before and after the pairing window (before the window: 15.3 ± 10.1; after the window: 25.9 ± 16.4, S1). This window coincided with the egg-laying peak (Figure 3B) and the dramatic decline in the number of single males, reflecting low mating opportunities for females (Figure 4C). Moreover, in the YRD population, males that paired up earlier produced a clutch sooner (S2). In the other two populations with a shorter breeding season, such a similar ‘efficient mating window’ was observed early in the season for the LHK population, where males spent on average only 6.6 days (n = 100 nests) finding a mate before day 80 (Figure 4B, 25.4 ± 10.7 days). However, this pattern was absent in the XH population, where males took an average of 16.5 days (n = 55 nests) to pair up with a female.

**Figure 4.**
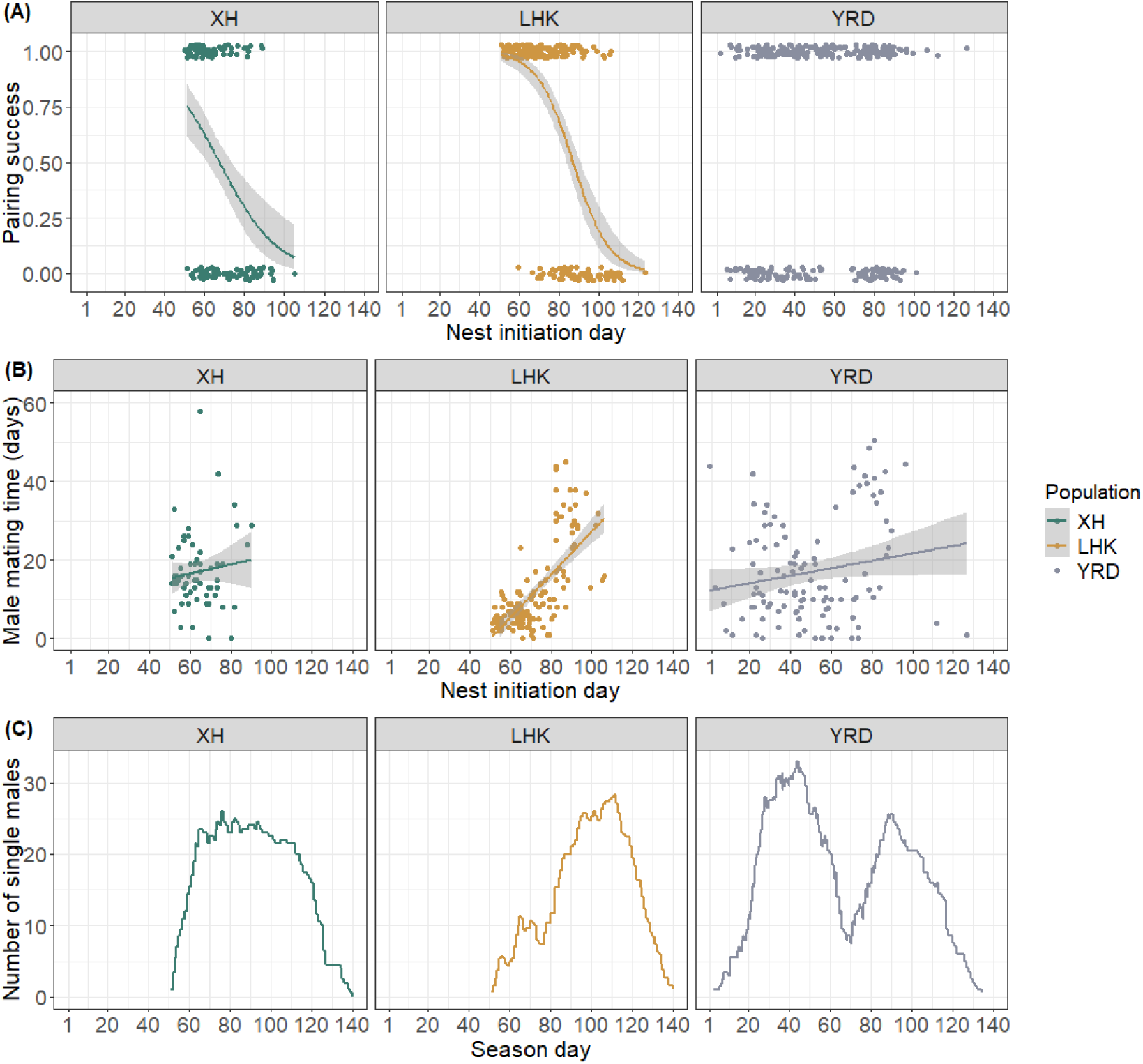
The seasonal trend of male (A & B) and female (C) mating opportunities in the three populations of Chinese penduline tits. (A) male pairing success (n_XH_ = 221 nests, n_LHK_= 277 nests, n_YRD_ = 391 nests; for statistical analyses see Table 3A); (B) mating time of males who paired successfully. Each dot in (A) and (B) represents nest initiation of a nest (n_LHK_= 141nests, n_XH_ = 61 nests, n_YRD_ = 113 nests; for statistical analyses see Table 3B). (C) the number of single males over the breeding season (n_XH_ = 109 nests, n_LHK_= 218 nests, n_YRD_ = 218 nests). X-axis: 1 = March 10^th^.

**Table 3.** Seasonal trends of male mating opportunities (i.e. pairing success and mating time) across the three populations of Chinese penduline tits (n_LHK_= 277 nests, n_XH_ = 221 nests, n_YRD_ = 391 nests)

| Response variable | Predictors | Chi-square ($\chi^2$) | df | P value |
| --- | --- | --- | --- | --- |
| (A) Pairing success | Nest initiation day | 20.96 | 1 | **<0.001** |
|  | Population | 20.33 | 2 | **<0.001** |
|  | Nest initiation day *<br>Population | 96.36 | 2 | **<0.001** |
| Seasonality of pairing success | XH | 19.61 | 1 | **<0.001** |
|  | LHK | 97.49 | 1 | **<0.001** |
|  | YRD | 0.22 | 1 | 0.64 |
| (B) Mating time | Nest initiation day | 262.78 | 1 | **<0.001** |
|  | Population | 308.31 | 2 | **<0.001** |
|  | Nest initiation day *<br>Population | 356.06 | 2 | **<0.001** |
| Seasonality of mating time | XH | 4.04 | 1 | **0.04** |
|  | LHK | 576.25 | 1 | **<0.001** |
|  | YRD | 33.72 | 1 | **<0.001** |

In all three populations, males that successfully paired spent significantly more time obtaining a mate as the season progressed (Figure 4b, Table 3b). In the LHK and XH populations, males were more likely to desert their nests when their mating opportunities were high and to stay for caring when mating opportunities were low. However, no such association was observed in the YRD population (Figure 5A). The flexibility of male parental care was evident as males adjusted their care decision over breeding attempts within a season in all three populations (S3).

**Figure 5.**
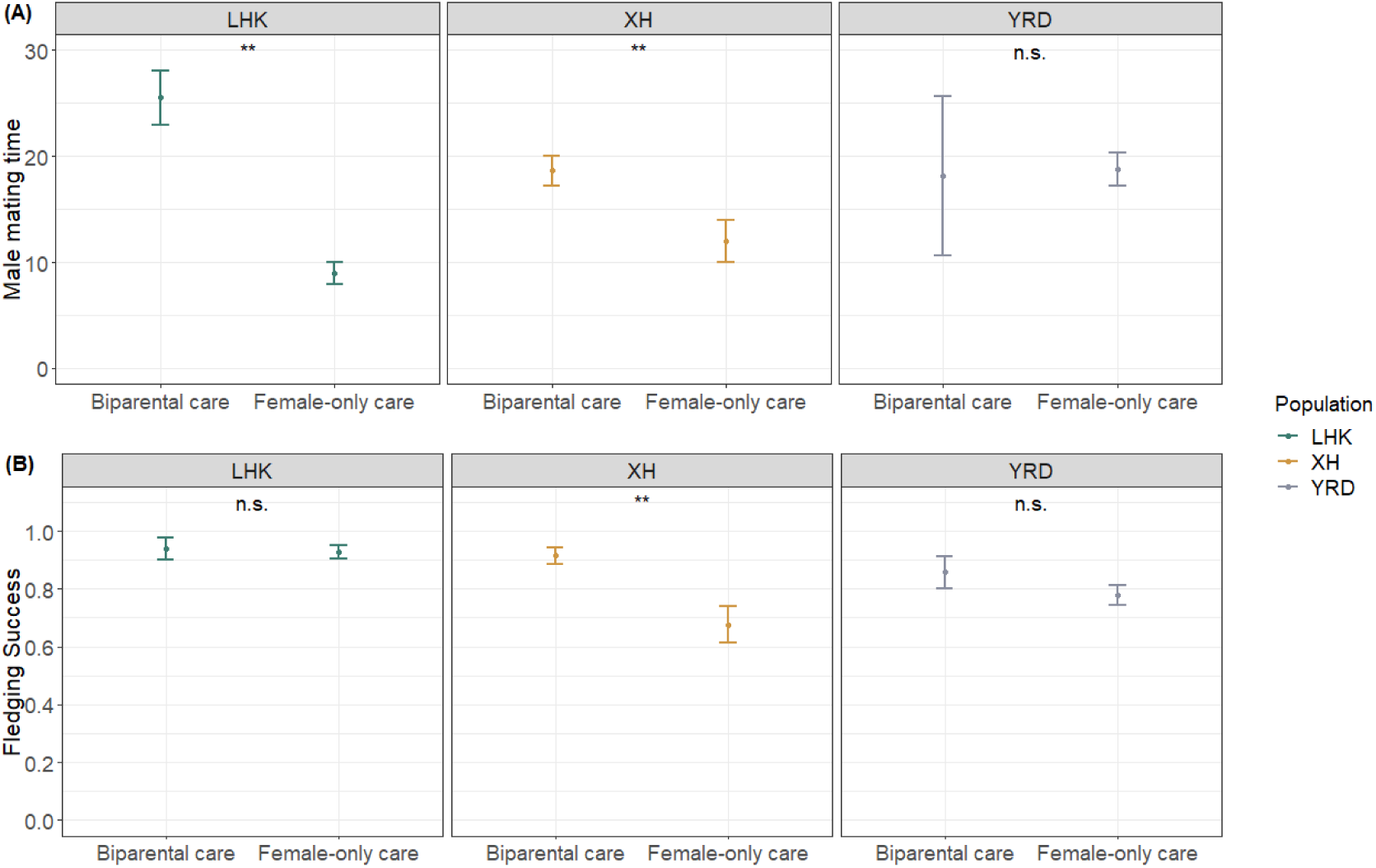
The associations of (A) male mating time and (B) fledging success with parental care in the three populations of Chinese penduline tits (male mating time: XH: *estimate*= 0.44, *se*= 0.09, *z*= 5.06, *p*< 0.001; LHK: *estimate*= 1.05, *se*= 0.06, *z*= 16.91, *p*< 0.001; YRD: *estimate*= 0.45, *se*= 0.04, *z*= 0.10, *p*= 0.99. Fledging success: XH: z= 4.60, p<0.01; LHK: z= 0.19, p= 0.85; YRD: z = 1.29, p=0.20).

Female mating opportunities increased throughout the season in the LHK and XH populations and decreased sharply toward the end of the season (Figure 4C). In the YRD population, female opportunities exhibited two peaks due to the occurrence of the ‘male efficient mating window’, during which most males deserted their first clutch and successfully remated. However, female care decisions were not influenced by mating opportunities in any of the three populations. Females consistently provided care throughout the season (XH: *t* = 0.30, *p* = 1.00; LHK: *t* = 0.30, *p*=0.99; YRD: *t* = 0.30, *p* = 1.00, n = 218 nests).

### Influence of nest desertion on fitness cost on the current broods

Average hatching success was similar in the LHK (0.82± 0.19, n= 54 nests) and XH (0.86 ± 0.20, n= 55 nests) populations, both of which were significantly higher than in the YRD population (0.77± 0.17, n = 93 nests, LHK vs XH--*z.ratio* = 0.71, *p*= 0.76; LHK vs YRD--*z* = 2.15, *p*= 0.08; XH vs YRD--*z*= 2.52, *p*=0.03). Hatching success did not differ among parental care patterns in any population (population*parental care: *χ^2^*= 3.46, *df*= 4, *p*=0.40). In contrast, fledging success varied significantly among the three populations (*χ^2^*= 19.51, *df*=2, *p<* 0.001), with the lowest success observed in the YRD population (Table 4, Figure 5B). In the XH population, fledging success was lower for uniparental care nests compared to biparental care nests, but such a difference was not found in the LHK and YRD populations (Table 4, Figure 5B).

**Table 4.** Fledging success of female-only care and biparental care nests in the three populations of Chinese penduline tits (n=152 nests)

| Population | Biparental care<br>(nests) | female-only care<br>(nests) | Chi-square ($\chi^2$) | p-value |
| --- | --- | --- | --- | --- |
| LHK | 0.94± 0.09 (8) | 0.92± 0.15(40) | 0.04 | 0.85 |
| XH | 0.91± 0.14 (26) | 0.72± 0.21 (19) | 22.63 | **<0.001** |
| YRD | 0.85± 0.17 (10) | 0.77± 0.24 (49) | 1.82 | 0.18 |

## Discussion

In this study, we revealed the complex parental care system in Chinese penduline tits, showing substantial variations both within and across populations: female-only care prevailed in the LHK and YRD populations, whereas biparental care is the main pattern in the XH population. Parental care varied seasonally in the LHK population, shifting from female-only care to biparental care, but no such trend was observed in the XH and YRD populations. Despite the YRD population having the longest breeding season, the peak timing of egg-laying was similar across all three populations. Our analyses suggest that interaction among mating opportunity, fledgling success and season length drove parental care variation across populations. In the LHK and XH populations, males were more likely to desert their clutch when mating opportunities were high but provided care when the opportunities were low. This association, however, was not found in the YRD population, which had a longer breeding season. Mating opportunity and its variation did not appear to influence female care decisions, as females mostly provided care across all three populations. In addition, we found that male desertion occurred more frequently in populations where fledging success did not differ between biparental and uniparental nests. In contrast, biparental care was the dominant strategy when fledging success became higher as a result of care from both parents compared to care from a single parent.

### Flexibility of parental care in relation to multiple local conditions

Our study revealed that parental care system in Chinese penduline tits is highly flexible and largely influenced by local conditions. Species with such variable parental care systems are generally rare but are widely distributed across taxa. For instance, neotropical poison frogs (family Dendrobatidae) switched from uniparental care to biparental care as a consequence of a reduced mating opportunity after their living space was experimentally reduced (Brown et al., 2008). In response to the threat of snake predation, uniparental care emerged in an introduced population of long-tailed sun skink (*Eutropis longicaudata*), a species that typically provides no care in the form of egg-protection (Pike et al., 2016). Biparental care is common in avian species, as offspring often rely heavily on close and considerable parental attendance to survive. This makes species, such as rock sparrow (*Petronia petronia*; Griggio & Pilastro, 2007; Griggio, 2015), barn owl (*Tyto alba*; Roulin, 2002; Béziers & Roulin, 2016), whiskered tern (*Chlidonias hybrida*; Ledwoń & Neubauer, 2017; Ledwoń et al., 2023), plovers (*Charadrius spp*.; Székely & Cuthill, 1999; Székely et al., 1999) and penduline tit (*Remiz spp.*; Ball et al., 2017; Zheng et al., 2021), which regularly exhibit variations in parental care valuable examples for studying the evolution of parental care flexibility.

While field experiments have examined the changes in population desertion rates in response to manipulated local conditions (i.e., providing food supply to change food abundance, Eldegard & Sonerud, 2009; capturing and releasing single males to manipulate mating opportunities, Székely et al., 1999), comparative analysis has approached the causes of parental care variations by revealing common associations between specific conditions (e.g., sex ratio or ambient temperature) and the distributions of care patterns across populations (Vincze et al., 2013; Eberhart-Phillips et al., 2018). For example, biparental care emerged more frequently under high-temperature constraints across ten geographic populations of plovers, as parents need to cooperate in incubating to stop eggs from being overheated (Vincze et al., 2013). This follows the same logic as meta-analyses that search for common associations between parental care and local conditions at the phylogeny levels (Remeš et al., 2015; Furness & Capellini, 2019; Vági et al., 2019; Vági et al., 2024). However, these studies found it difficult to illustrate how the associated factors substantially shape parental care variations. By systematically comparing the breeding phenology, dynamics of local conditions, and the seasonal trends of parental care across three populations of Chinese penduline tits, our study suggests that the formation of parental care variations in nature is more complex than simply identifying direct associations with one factor at a time. These variations require more careful explanations.

### Males responded to mating opportunities that varied differently across populations

We found that similar parental care patterns in Chinese penduline tits between populations LHK and YRD were not caused by the same reasons. In the LHK population, males collectively deserted their clutch early in the season when mating opportunities were high, and switched to provide care as these opportunities declined in the late season. In contrast, the situation was more complex in the YRD population. Despite relatively low mating opportunities in the early season, males in YRD still predominantly deserted their first clutch just before an ‘efficient mating window’, during which days most males quickly acquired a female partner and then sought a new mate. This indicates that male penduline tits can quickly sense the temporal state of local conditions (such as the increased availability of unmated females) and adjust their parental decision accordingly. This flexible reaction in parental care was recorded across wide taxa, for example, male cichlid fishes are more likely to terminate care when their mating opportunity is high (Balshine-Earn & Earn, 1998); female snowy plovers (*Charadrius nivosus*) were more likely to desert the brood when they sense higher mating opportunity within the season (Kupán et al., 2021); and in dendrobatid frog (*Allobates femoralis*), female deserters resumed tadpole transport upon sensing that their male partners has been experimentally removed in wild (Ringler et al., 2015). The collective brood desertion of male penduline tits in the YRD population around the ‘efficient mating window’ indicates that the perceived mating opportunities in upcoming days may play a more important role in shaping male care decisions than the opportunities they experienced before producing the current brood. This aligned with the finding in burying beetles (*Nicrophorus vespilloides*), where males extended their care duration in response to high male mate competition in the group (Hopwood et al., 2015).

In addition, we observed that females likely controlled the pace of breeding throughout the season. In the YRD population, females appeared less imperative in entering breeding status during the early season, or they arrived at the breeding grounds later in batches than males. The closely timed egg-laying peaks in the three populations reflected that females scheduled their breeding by tracing the ecological phenology rather than the timing of mate acquisition. The rapid increase of available unmated females around day 50 likely shaped the ‘efficient mating window’ for males in the YRD population. Nevertheless, the question remains why many males initiated mate attraction much earlier than the breeding peak in the YRD population. We found that males who paired up early produced their clutch early. The long breeding season provided early-mated males with additional benefits, as they could still catch up with the ‘efficient mating window’ after desertion, allowing them to produce another clutch before the peak of female egg-laying. This trend was not observed in the short-seasoned LHK and XH populations, where males faced more intense time constraints to breed.

The effects of season length have received relatively little attention in previous studies of parental care variations. For example, by temporarily removing nest boxes in the early season to simulate a shortened breeding season, Griggio (2015) found that rock sparrows (*Petronia petronia*) in the experimental area showed a higher rate of offspring desertion compared to those in the control area, where nest boxes remained available. However, a contrasting conclusion came from a mate-analysis showing that shorter fruiting season lengths resulted in less male desertion in frugivores (Barve & La Sorte, 2016). One theoretical model resembling the penduline tit system has suggested that the effect of season length may be related to the duration of a single breeding cycle (Zheng et al., 2023). Our results demonstrate that the influence of breeding season on parental care is complex. It does not simply affect the maximum number of breeding attempts birds can fit into the season, but also interacts with other factors, such as ecological phenology and mating opportunities, as we have demonstrated here. These interactions collectively shape the parental care strategies of animals.

### Fitness costs of offspring desertion matter for making parental care decisions

These complex effects from multiple factors also existed in the XH population, where male parental care was contrastingly associated with different local conditions. Although male penduline tits in the XH population also faced seasonally decreased mating opportunities, which would typically motivate them to desert their first clutch, instead, most males did not desert but provided care to their brood. This can be explained by the significantly lower fledging success in uniparental care nests compared to biparental care nests in this population. In other words, offspring desertion would entail fitness costs for the current brood, which reduced the propensity of male desertion. Conversely, offspring desertion did not result in any fitness costs in the other two populations, where the uniparental and biparental care nests had similar fledging success. The absence of fitness costs may be affected along with the high mating opportunities, likely escalating the propensity of male desertion in the LHK and YRD populations, as they could effectively free-ride on their partner’s parental efforts. This suggests that male penduline tits made care decisions by balancing reproductive payoffs. This aligns with findings in snowy plover, where offspring desertion occurs only when a partner is competent to raise the rest of the brood alone (Kupán et al., 2021). Similarly, female Tengmalm’s owl (*Aegolius funereus*) desert the brood only in good years when the male partner can collect sufficient resources to raise the nestlings alone (Eldegard & Sonerud, 2009).

Uniparental care has generally been thought to evolve only when single parents can raise offspring with high efficiency (Székely et al., 1996; Roulin, 2002; Webb et al., 2002; Klug & Bonsall, 2010; Kupán et al., 2021). However, our study shows that this view does not entirely apply to Chinese penduline tits. Although uniparental care predominated in the YRD population, the reproductive success of uniparental nests in this population was surprisingly significantly lower than that of uniparental care nests in XH, a biparental care dominated population. This outcome offers a new insight that challenges previous understanding: the difference in reproductive success between uniparental and biparental care nests may be more influential in the evolution of parental care than the absolute reproductive success of uniparental care, because the former reflects the additional fitness costs resulting from offspring desertion.

### Female penduline tits do not respond to local conditions in parental care

Our study found that female Chinese penduline tits consistently provided care regardless of the local conditions across all three populations. Female desertion was relatively rare, and females did not respond to variations in mating opportunity. This result contrasts with several previous studies suggesting that the rarer sex is more likely to desert offspring to remate and increase fitness when mate availability is high (Székely et al. 2014, Schacht & Borgerhoff Mulder, 2015; Rosa et al., 2017; Eberhart-Phillips et al., 2018; Henshaw et al., 2019). We found that females penduline tits adhered closely to phenological cues and employed a highly consistent parental care strategy, which strikingly contrasts with the flexible male strategy in breeding in each of the three populations. Such sex-different responses in parental care have often been demonstrated by mate removal experiments. For example, after mate removal, male burying beetles significantly increased their care level, while females showed no response to the partner’s absence (Suzuki & Nagano, 2009). Similarly, during short-term mate removal experiments, male rock sparrows compensated for provisioning frequency, but females showed no change (Cantarero et al., 2019). However, recent research on Chinese penduline tits revealed that in biparental care nests, both males and females fully compensated for provisioning when their partner was removed (Zheng et al., 2024). Therefore, parental responses to external conditions and changes in a partner’s care effort may evolve under different mechanisms. For instance, the sex with higher annual mortality rates may cherish more of the current reproduction, therefore, stay more frequently for caring (Klug et al., 2013; Henshaw et al., 2019); sex divergent physiological condition (such as hormone level of prolactin or testosteron) may induce one sex to care more than the other (Hunt et al., 1999; Smiley, 2019); frequency-dependent selection acts through sex-divergent dispersal rate may shape strategies of breeding behaviour (Schwanz & Georges, 2021). Factors such as extra-pair paternity have also been found to influence parental care in animals, as they deflated male fitness and may reduce male care for offspring (Griffin et al., 2013; Ball et al., 2017). However, the three Chinese penduline tits populations showed similar proportions and a low level of extra-paired paternity (in 10-20% nests, unpublished data), suggesting that this may not be a factor causing the variation in parental care observed in this species.

### Conclusions and recommendations for future research

Our study revealed a highly flexible parental care system of Chinese penuline tits and demonstrated sex-divergent parental care strategies in response to local conditions. Male parental care decision was significantly influenced by seasonally varied fitness payoffs shaped by season length, reproductive success and mating opportunities. Females, however, showed no response to local conditions when making parental care decisions. Our study emphasis the importance of considering comprehensive environmental effects in the evolution of parental care by carrying out systematic investigations across populations in wild species. Currently, there is limited knowledge about the causes behind the differing fledging success between care types and across populations. Further quantitative analysis of environmental conditions (e.g., food abundance, parasite prevalence) and physiological conditions of both sexes, as well as provisioning frequency across populations, will be promising for clarifying these patterns. Future studies should integrate sex-specific life-history traits and biological conditions to better understand the evolution of sex-divergent responses in parental care.

