## supplement for "Flexible parental care in a songbird correlates with sex-specific responses to seasonal phenology, mating opportunity and reproductive success"

This Supplement includes:

**Supplement Figure 1.** Illustration of the efficient mating window (days 55-70) for male Chinese penduline tits in the YRD population.

**Supplementary Figure 2.** Positive correlation between male pairing-up day and start egg-laying day for Chinese penduline tits in the YRD population.

**Supplementary Figure 3.** Flexibility of male parental care decisions across breeding attempts within a breeding season in each of the three Chinese penduline tit populations.

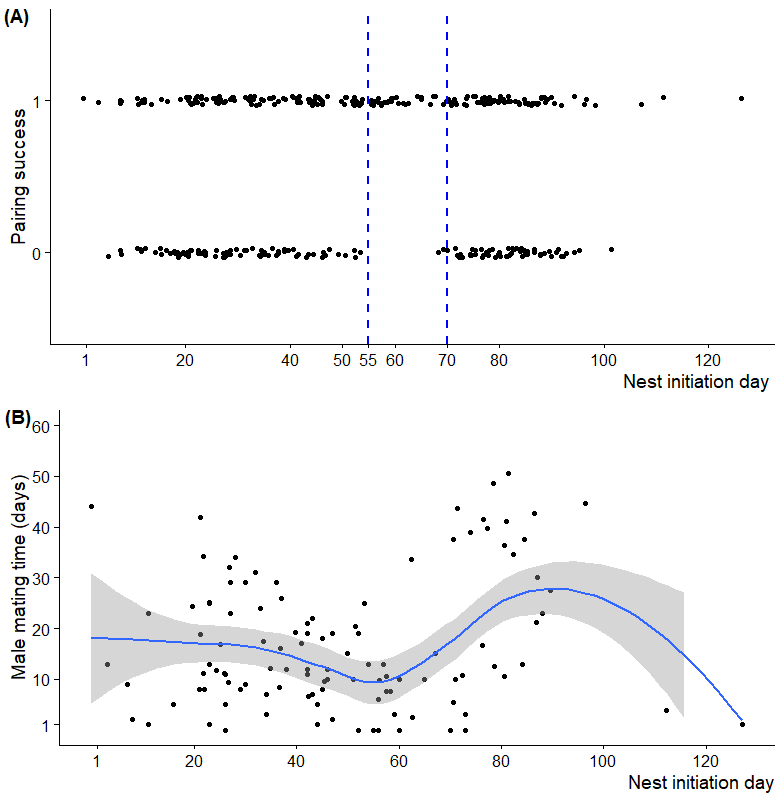

**Supplementary Figure 1.** Illustration of the efficient mating window (days 55-70) for male Chinese penduline tits in the YRD population. (A) shows the high pairing up success for nests initiated during this period. The two blue dashed lines indicate the position of the quick mating window, almost all the males with nests succeeded in finding a mate during this time (89%; 24 out of 27 males). (B) shows the variation of male mating time over the season. Blue and black dots indicate nests succeeded or failed in pairing up with a female. The smoother depicts the seasonal trend of mating time of males over the season and the gray ribbon indicates the 95% confidence level interval.

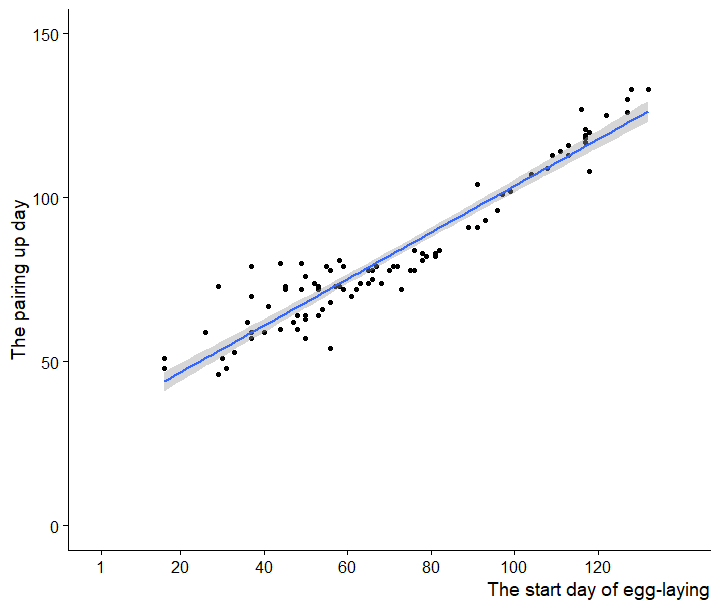

**Supplementary Figure 2.** Positive correlation between male pairing-up day and start egg-laying day for Chinese penduline tits in the YRD population (*χ^2^*= 1064.5, p< 0.001, n = 134 nests). Each dot indicates a nest and the blue line manifested the relationship of the two variables by predictions from the linear regressions, with gray ribbons indicating the 95% confidence level interval.

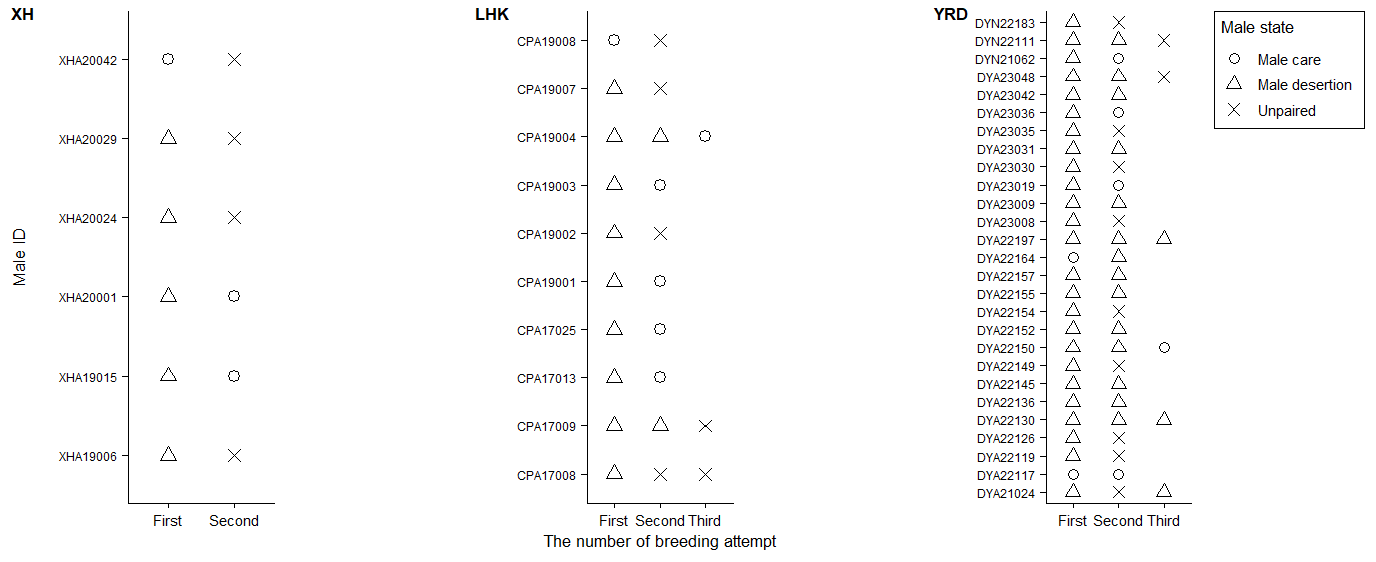
**Supplementary Figure 3.** Flexibility of male parental care decisions across breeding attempts within a breeding season in each of the three Chinese penduline tit populations. The Y-axis represents males successfully tracked over multiple breeding attempts, with different symbols indicating their parental care decisions or pairing status in each attempt. “Male care” includes males in biparental care and male-only care nests, while “male desertion” includes males in female-only care and biparental desertion nests. “Unpaired” denotes males that failed to acquire a new mate after brood desertion.
